# The parallel-stranded d(CGA) duplex is a highly predictable structural motif with two conformationally distinct strands

**DOI:** 10.1101/2021.09.30.462591

**Authors:** Emily M. Luteran, Paul J. Paukstelis

**Affiliations:** Department of Chemistry and Biochemistry, University of Maryland, College Park, MD, 20742, USA

## Abstract

DNA can adopt non-canonical structures that have important biological functions while also providing structural diversity for nanotechnology applications. Here, we describe the crystal structures of two oligonucleotides composed of d(CGA) triplet repeats in the parallel-stranded duplex form. The structure determination of four unique d(CGA)-based parallel-stranded duplexes across two crystal structures has allowed us to characterize and establish structural parameters of d(CGA) triplets in the parallel-stranded duplex form. Our results show that d(CGA) units are highly uniform, but that each strand in the duplex is structurally unique and has a distinct role in accommodating structural asymmetries induced by the C-CH+ base pair.

## INTRODUCTION

DNA is a polymorphic biopolymer that can adopt an array of conformations beyond the traditional B-form double helix. Watson-Crick base paired duplexes can access multiple helical forms (A or Z-form) depending on sequence and environmental conditions (1,2). The hydrogen bonding and base stacking interactions that stabilize anti-parallel duplexes also allow DNA to access other non-canonical conformations, some of which have known biological implications, including G-quadruplexes (3), i-motifs (4), and triplexes (5). Further, the formation of many non-canonical structures can be controlled by nucleotide sequence composition and several environmental factors including pH, the presence and concentration of cations, and temperature (6–11). The alternative structures formed by genomic DNA triplet repeat sequences (12–14) have been implicated in their ability to expand and cause genetic instabilities (15,16). Understanding these alternative structures and the conditions that may lead to their formation provides a fundamental basis for understanding the disease states. Additionally, the ability to control DNA conformations has utility in DNA nanotechnology applications where non-canonical motifs expand structural and functional diversity of nanostructures while retaining inherent programmability and predictability. Previous work has focused on incorporating functional non-canonical structures such as the G-quadruplex (17–20), i-motif (21–24), polyA-motif (25–27), and triple helix (28–30) into DNA nanostructures, where changes in the local environment are used to tune the resulting structure.

The d(CGA) triplet repeat motif is another such environmentally sensitive motif that can adopt different structural forms in a pH-dependent manner at near-physiological temperature and salt concentration (31). Neutral pH favors a unimolecular anti-parallel hairpin stabilized by canonical G-C base pairs (12,32), while acidic pH favors a non-canonical homo-base paired parallel-stranded duplex (ps-duplex) (33,34). Although the non-canonical d(CGA) motif can adopt distinct structural conformations, the ps-duplex is the predominantly studied form (33,35–38). Originally described as **Π**-DNA, the d(CGA)_n_ ps-duplex is stabilized by homo-base pair interactions (C-CH+, G-G, and A-A) and inter-strand base stacking interactions (34,38). The C-CH+ homo-base pair requires hemi-protonation at the N3 position to form three hydrogen bonds along the Watson-Crick face (34). N2-N3 sugar-edge hydrogen bonds stabilize G-G homo-base pairs, while A-A homo-base pairs are formed through N6-N7 Hoogsteen face hydrogen bonds. Importantly, the GpA dinucleotide step provides significant stabilization to the ps-duplex by the formation of interstrand G/A base stacking interactions.

The structure and stability of the ps-duplex is highly influenced by the 5ʹ-nucleotide of each triplet (31). Similar G/A-stacking interactions have been observed in ps-duplex structures containing d(GGA) or d(TGA) triplets (35,37,39–41), though contiguous repeats of these sequences are unable to form ps-duplexes (31). The ps-duplex region of an intercalation-locked tetraplex containing d(TGA) triplets forms a perfectly symmetrical duplex (37), while the same ps-duplex region containing d(CGA) triplets resulted in structural asymmetry and duplex bending (36). The asymmetry is associated with a displacement from the helical axis at the C-CH+ base pair. Further, thermodynamic studies indicated that ps-duplex structures containing six tandem d(YGA) triplet repeats undergo a significant destabilization when d(CGA) triplets are replaced with d(TGA) triplets (31). Beyond the additional hydrogen bond interaction within each C-CH+ base pair, the structural details as to why asymmetric d(CGA) duplexes are significantly more stable than symmetric d(TGA) triplets remain unclear.

The ability of d(CGA) to form distinct structural states appears to be a trait shared by several other triplet repeat motifs, though d(CGA) is the only triplet known to form perfectly ps-duplex structures (12–14,31). Interestingly, genomic analysis of all possible triplet repeat sequences have shown that d(CAG) triplets are over-represented in the human genome and indicated in disease pathologies while d(CGA) triplet repeat sequence are the least frequently observed, occurring only 16 times (42). A similarly low frequency and coverage of d(CGA) triplets was seen when a comparable genomic analysis was performed in other eukaryotic organisms (43). The formation of alternative structures and the relative stabilities of such structures are thought to be important factors contributing to the expansion of repeat sequences (13,44,45). Due to the challenges they present to replication machinery, repeat sequences that form alternative structures could influence pathological or evolutionary outcomes (32,46). Therefore, it is important to characterize the structural diversity of triplet repeat sequences that have the ability to form such non-canonical structures.

In this work we have determined the crystal structure of two oligonucleotides containing multiple tandem d(CGA) triplet repeats in the ps-duplex form. These structures are the longest ps-duplexes to be solved solely comprised of such triplets. The crystals grew from different solution conditions and resulted in distinct crystal packing arrangements. The structure determination of four ps-duplexes across these two different crystal structures has allowed us to thoroughly characterize and define the structural features of d(CGA) triplets and the ps-duplexes they form. Despite crystallization and molecular packing differences, the resulting ps-duplex structures have strikingly low RMSD values, demonstrating the robust structural uniformity of the d(CGA) triplet repeat motif in the ps-duplex form. Additionally, each ps-duplex contains two conformationally distinct d(CGA) triplets based on hydrogen bonding and base stacking interactions. Surprisingly, each strand contains only one triplet conformation. Thus, ps-duplexes containing d(CGA) repeats are not structurally symmetrical and the apparent structural asymmetry is propagated discretely throughout each strand.

## MATERIAL AND METHODS

### Oligonucleotide Synthesis and Purification

DNA oligonucleotides were synthesized on the 1 μmol scale using standard phosphoramidite chemistry on an Expedite 8909 Nucleic Acid Synthesizer (PerSeptive Biosystems, Framingham, MA) with reagents from Glen Research (Sterling, VA). Following deprotection with 30% ammonium hydroxide, oligonucleotides were purified by 20% (19:1) acrylamide/bis-acrylamide, 7 M urea gel electrophoresis, electroeluted, and dialyzed against deionized water.

### Oligonucleotide Crystallization

(CGA)_5_TGA was crystallized by mixing 2 μL of 200 μM DNA solution with 2 μL of crystallization solution (20% 2-methyl-2,4-pentanediol (MPD), 120 mM barium chloride, 30 mM sodium cacodylate, pH 5.5). GA(CGA)_5_ was crystallized by mixing 1 μL of 125 μM DNA solution with 2 μL of crystallization solution (8% PEG400, 96 mM strontium chloride, 32 mM lithium chloride, 8 mM hexamminecobalt(III) chloride, 24 mM sodium cacodylate, pH 7.4). Crystallization was performed in sitting drops, equilibrated against 300 μL of 30% MPD or PEG400 (for (CGA)_5_TGA and GA(CGA)_5_, respectively) in the well reservoir, and incubated at 22°C. Crystals were observed within 7 days of plating.

### Data collection, processing, and structure determination

Crystals were removed from drops with nylon cryo-loops, immediately dipped in the respective crystallization condition supplemented with 30% MPD or PEG400 and plunged into liquid nitrogen. Diffraction data were collected at the Advanced Photon Source (APS), Argonne National Laboratory. (CGA)_5_TGA data was collected on the 24-ID-E beamline and GA(CGA)_5_ data was collected on the 24-ID-C beamline.

Data processing for (CGA)_5_TGA was carried out with iMosflm (47) and GA(CGA)_5_ was carried out with XDS (48) and Aimless (49). Initial phases were obtained by molecular replacement using Phaser (50). The parallel stranded homo-duplex d(CGA) triplet region from PDB id: 1IXJ (35) was used as the search model for (CGA)_5_TGA, and two tandem d(CGA) units from the refined (CGA)_5_TGA structure were used as the search model for GA(CGA)_5_. Model building and refinement was carried out for in Phenix (51) and Coot (52), respectively, for both datasets.

### Circular Dichroism

CD spectra were obtained using a Jasco J-810 spectropolarimeter fitted with a thermostatted cell holder (Jasco, Easton, MD). Samples were prepared using 10 μM DNA in 20 mM MES, 100 mM sodium chloride (pH 5.5), or 20 mM sodium cacodylate, 100 mM sodium chloride (pH 7.0). Samples were incubated at 4 °C overnight prior to data collection. Data were collected at room temperature at wavelengths from 220-300 nm.

## RESULTS AND DISCUSSION

### Overview

We determined the crystal structures of (CGA)_5_TGA and GA(CGA)_5_ in the ps-duplex form at 2.1 Å and 1.32 Å, respectively (Table 1). (CGA)_5_TGA was crystallized at pH 5.5 to preferentially stabilize the ps-duplex form, while GA(CGA)_5_ was crystallized at pH 7.4 to characterize the anti-parallel hairpin form. Despite being at a pH that strongly favors the hairpin form (Figure 1A,B), GA(CGA)_5_ also crystallized as a ps-duplex. Several other structures that rely on the C-CH+ hemi-protonation have also crystallized as ps-duplexes at above-neutral pH, suggesting that factors beyond pH influence this structural preference (37,54). The high local concentration of DNA and the presence of crowding agents have been demonstrated to increase the observed pH of the structural transition in C-CH+ mediated structures (32,55,56). CD measurements of d(CGA) repeat sequences are consistent with these observations; the presence of crowding agents shifts the favorability range of the ps-duplex to higher pH (Figure 1C). Specifically, the addition of 30% PEG2000 increased the pH of the structural transition by 0.33 ± 0.06 and 0.32 ± 0.13 pH units for (CGA)_5_TGA and (CGA)_5_, respectively (Figure 1D). Also, previous thermodynamic measurements have demonstrated a significantly greater stability in the ps-duplex over the anti-parallel hairpin form (31). Therefore, it is not surprising that the significantly more stable ps-duplex form is dominant in crowded crystallization conditions where structural stability is advantageous. It may also thus be possible for d(CGA) ps-duplexes to form in crowded cellular environments, similar to other C-CH+ mediated DNA structures (57–59). Despite testing multiple constructs of d(CGA)-derived oligonucleotides, we were unable to determine a structure in the hairpin form.

**Table 1.**
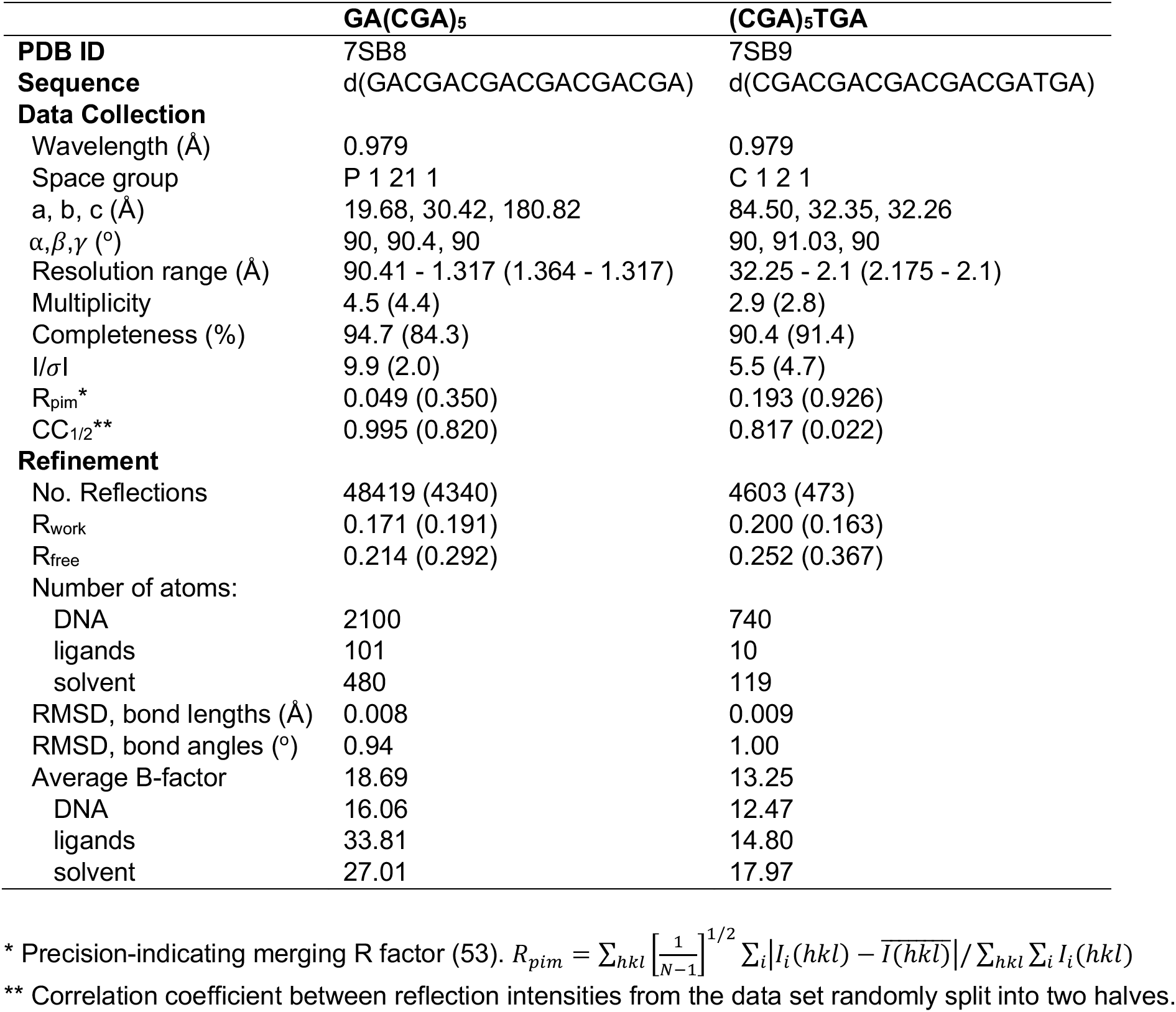
Data collection and refinement statistics. Values in parentheses correspond to the high-resolution shell.

**Figure 1.**
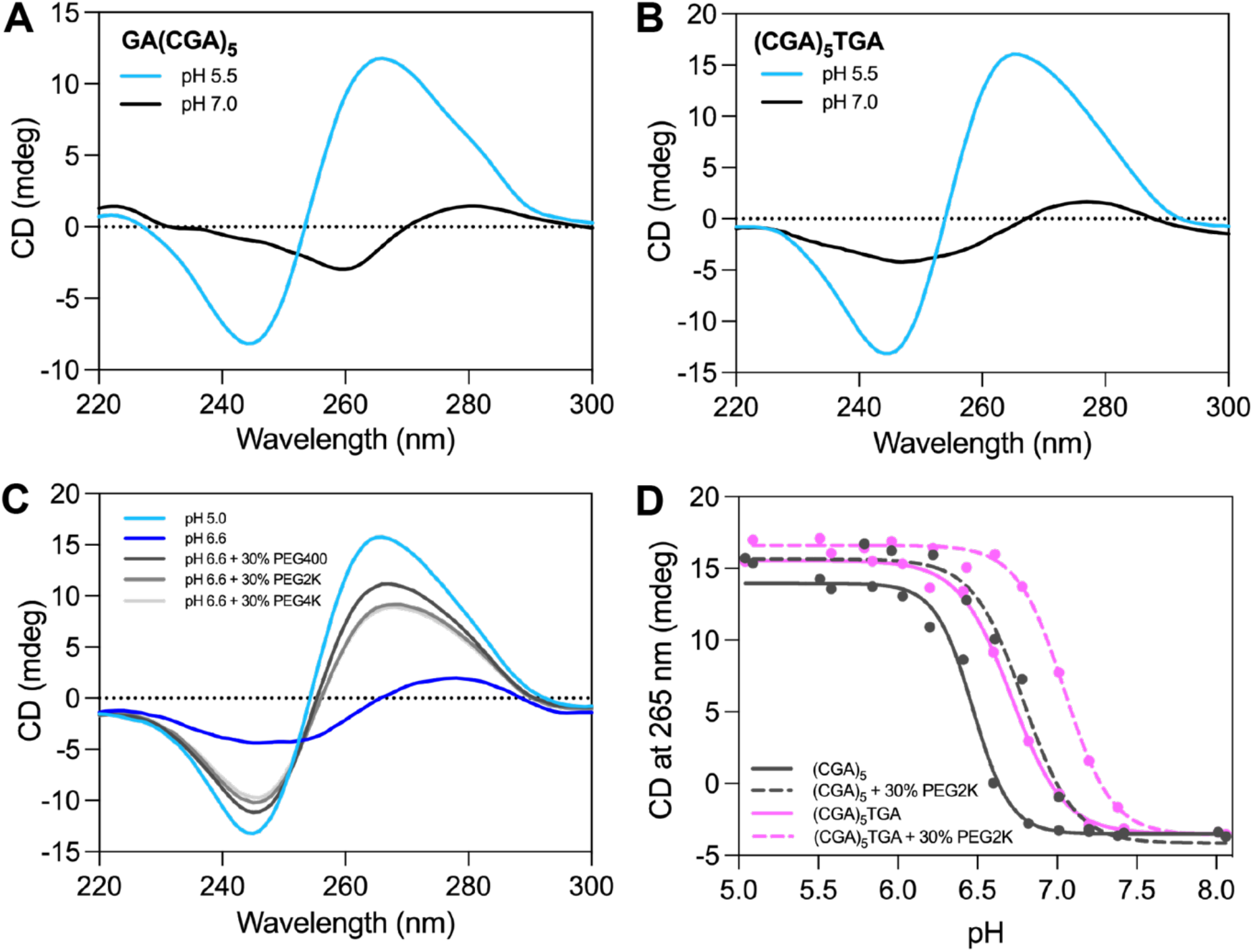
CD spectra of d(CGA)-containing oligonucleotides. The positive band at 265 nm and negative band at 245 nm are characteristic of the ps-duplex form (31,32,34). The anti-parallel form has a weak positive band at 280 nm and weak negative band at 260 nm (31,32,34). **A**. CD spectrum for GA(CGA)_5_ at pH 5.5 (blue) or pH 7.0 (black). **B**. CD spectrum for (CGA)_5_TGA at pH 5.5 (blue) or pH 7.0 (black). **C**. (CGA)_5_ forms a ps-duplex at pH 5.0 (light blue) and anti-parallel hairpin at pH 6.6 (dark blue) in the same buffer conditions as A/B. The formation of the ps-duplex form at pH 6.6 is favored in the presence of 30% PEG400 (dark gray), PEG2000 (medium gray), and PEG4000 (light gray). **D**. Crowding agents increase the pH of the structural transition from ps-duplex to anti-parallel hairpin form. The transition was measured as the loss of characteristic ps-duplex signal at 265 nm in native conditions (solid lines) or in the presence of 30% PEG2000 (dashed lines) for (CGA)_5_ (gray) and (CGA)_5_TGA (pink).

### Crystal Packing

In the GA(CGA)_5_ crystal structure, six strands form three parallel-stranded homo-duplexes (Duplex 1, 2, and 3) in the asymmetric unit (Figure 2A). Duplex 2 is coaxially stacked between duplexes 1 and 3 though 3ʹ to 5ʹ end stacking of the terminal G1-G1 and A17-A17 base pairs. This arrangement results in a junction of three tandem sets of interstrand G/A stacking interactions at each duplex intersection to stabilize the crystal lattice (Figure 2B). This packing arrangement forms columns of alternating ps-duplexes propagating throughout the crystal along the c-axis. Interestingly, this is the first instance of 3ʹ to 5ʹ end stacking in this class of ps-duplexes; other ps-duplexes containing the d(CGA) motif stack in the 3ʹ-3ʹ or 5ʹ-5ʹ orientation (36). This difference is likely due to the lack of 5ʹ-C. The exposed 5ʹ-G allows for preferential formation of inter-duplex G/A stacking interactions with the 3ʹ-A of another duplex that directly mimic the internal interstrand G/A stacking interactions. In the (CGA)_5_TGA structure, two strands form one homo-duplex (Duplex 4) in the asymmetric unit (Figure 2C). The duplex is stacked with crystallography identical duplexes via 5ʹ-5ʹ stacking of C1-C1 base pairs and 3ʹ-3ʹ stacking of A18-A18 base pairs.

**Figure 2.**
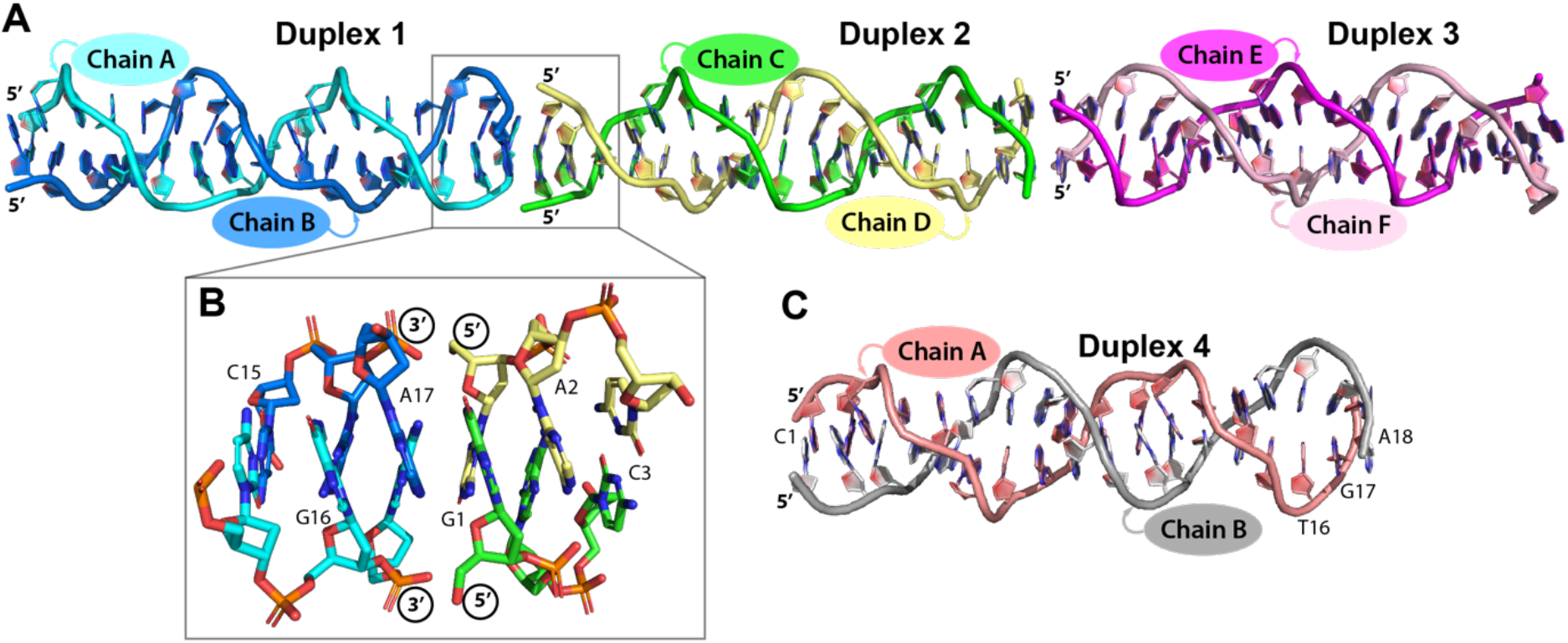
Overview of the d(CGA)-based parallel-stranded homo-duplexes. **A**. The asymmetric unit for GA(CGA)_5_. The individual chains within each duplex (1–3) are labeled and colored accordingly. Duplex 1: chain A (cyan), chain B (blue); Duplex 2: chain C (green), chain D (yellow); Duplex 3: chain E (magenta), chain F (light pink). **B**. 3ʹ to 5ʹ end stacking of duplex 1 and 2. The 3ʹ A17-A17 base pair of duplex 1 forms stacking interactions with the 5ʹ G1-G1 base pair of duplex 2 to form three tandem G/A stacking interactions. **C**. The (CGA)_5_TGA asymmetric unit. Each chain within duplex 4 is labeled and colored accordingly: chain A (salmon), chain B (gray).

Both crystals grew in the presence of divalent cations where they primarily mediate inter-duplex crystal packing interactions (Figure S1). When possible, anomalous difference maps and coordination distances were used to verify cation identity and placement (Figure S2). GA(CGA)_5_ (duplexes 1-3) crystallized in the presence of hexamminecobalt(III) (NCO) and Sr^2+^. Specifically, NCO is positioned in multiple conformations between the Hoogsteen faces of guanosines from two duplexes, with GN_7_-GN_7_ and GO_6_-GO_6_ distances of 8.7 ± 0.3 Å and 7.6 ± 0.1 Å, respectively (Figure S1A). Sr^2+^ mediates the remaining inter-duplex guanosine positions in two distinct modes. The first set of Sr^2+^ mediated interactions are similar to NCO but have shorter GN_7_-GN_7_ and GO_6_-GO_6_ distances (7.9 ± 0.1 Å and 6.8 ± 0.1 Å, respectively) (Figure S1B). The remaining Sr^2+^ cations are similarly positioned between two guanosines from separate ps-duplexes but the major groove faces are positioned such that GN_7_-GO_6_ are oriented together (9.31 ± 0.03 Å) (Figure S1C). In the (CGA)_5_TGA structure, Ba^2+^ mediates inter-duplex packing in two distinct environments. One mode is almost identical to the first set of Sr^2+^ mediated interactions, where GN_7_-GN_7_ and GO_6_-GO_6_ distances are 7.8 ± 0.1 Å and 6.9 ± 0.0 Å, respectively (Figure S1D). The remaining Ba^2+^ cations are positioned between the major groove face of one guanosine and the phosphate oxygen of the opposing duplex guanosine where the GN_7_-PO_2_ and GN_6_-PO_2_ distances are 9.4 ± 1.3 Å and 8.9 ± 0.9 Å (Figure S1E). Interestingly, despite the different cations with unique packing interactions, the resulting ps-duplex structures were highly uniform.

### d(CGA) ps-duplexes are structurally isomorphous and highly uniform

Though these structures were solved from individual crystals with different DNA sequences, solution conditions, and crystal packing arrangements, the resulting ps-duplex structures are nearly identical over the length of tandem d(CGA) repeats (Figure 3A). The three duplexes from the GA(CGA)_5_ structure have RMSD values between 0.421 Å and 0.451 Å for 700 atoms fit (Figure 3B). Most of the structural deviation arises from subtle differences in the phosphate backbones, likely resulting from solvent interactions that influence crystal packing (Figure 3C). Despite being crystallized in different conditions and containing the C16T substitution, duplex 4 is also highly similar to duplexes 1-3 (respective RMSD values: 0.846 Å, 0.855 Å, 0.877 Å for 698 atoms fit) (Figure 3A,B). The structural deviations associated with duplex 4 are primarily observed near the substitution site. Weaker electron density and correspondingly higher B-factors observed from A12 to A18 in duplex 4 may also contribute to increased RMSD values, though the overall ps-duplex structure is maintained.

**Figure 3.**
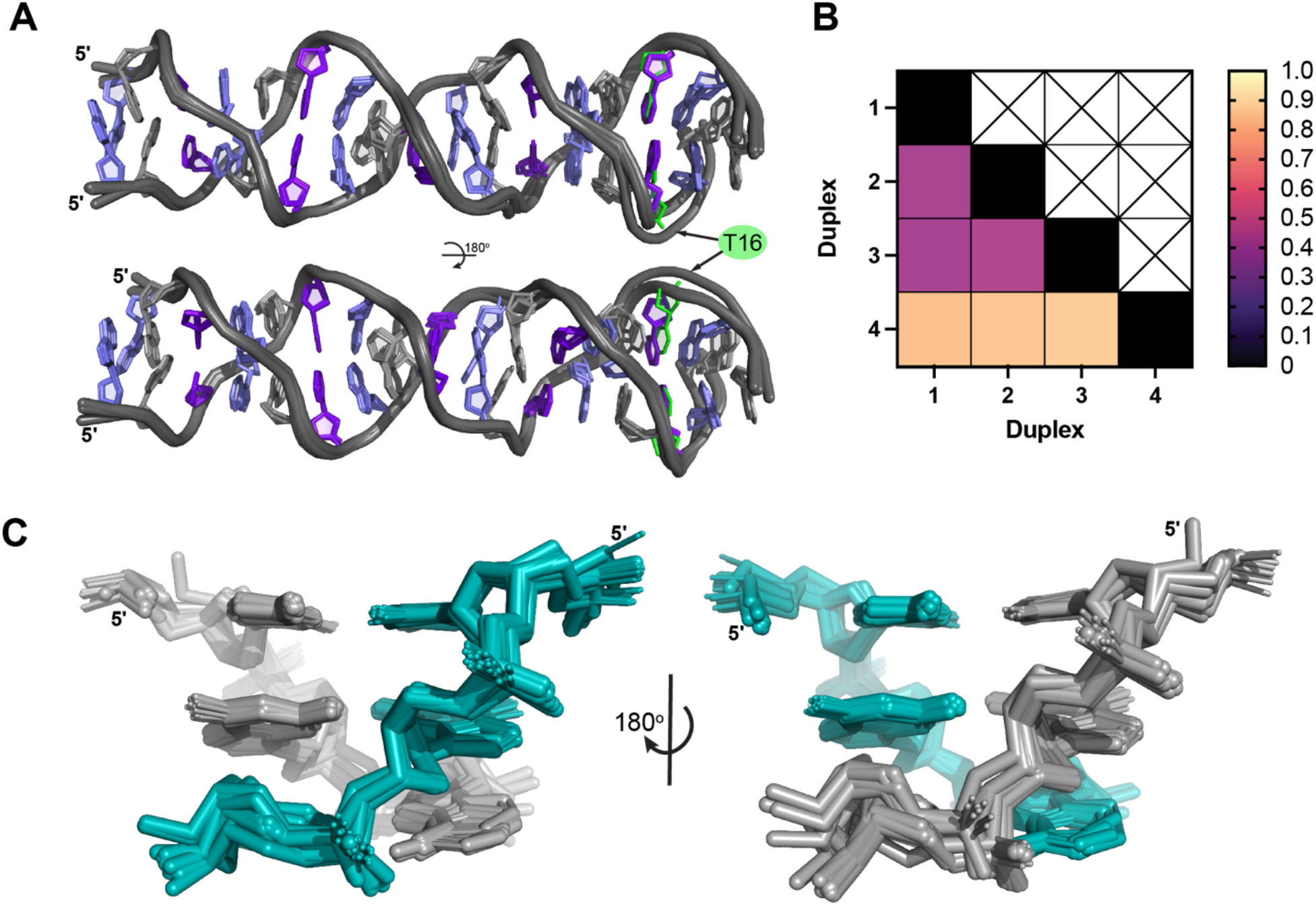
d(CGA) triplets in the ps-duplex form are structurally isomorphous. **A**. Overlay of duplexes 1-4 illustrates the robust structural uniformity of the ps-duplex form across different sequences, solution conditions, and crystal packing arrangements. Structural deviations are primarily observed surrounding the C16T substitution position in (CGA)_5_TGA. Nucleotides are colored as follows: C (purple), G (light purple), A (gray), T (green). The 5’ C-CH+ homo-base pair was omitted from the (CGA)_5_TGA in this overlay for simplicity. **B**. RMSD values from pair-wise alignment of duplexes 1-4. All ps-duplexes are highly similar with RMSD values below 1.0. **C**. Overlay of all d(CGA) triplets from duplexes 1-4 rotated 180° to show the difference in deviation along the phosphate backbone from each strand (colored teal or gray). Minimal overall deviations demonstrate the high structural predictability of the ps-duplex form of the d(CGA) triplet.

We also compared structures of isolated d(CGA) base paired triplets from all three duplexes to triplets within each duplex (intra-duplex) or from other duplexes (inter-duplex). Not surprisingly, comparison of all individual d(CGA) triplets results in high similarity as evident by the low RMSD of the full duplexes. Individual d(CGA) triplets at different positions in the same duplex (intra-duplex d(CGA) triplets) are almost identical (0.122 Å to 0.557 Å for 124 atoms fit) (Figure S3), indicating that there are no position-specific structural features along the helical length. The 3ʹ-end d(CGA) triplet of duplexes 1-3 and 3ʹ-most d(CGA) triplet of duplex 4 are the sources of the largest deviations among intra-duplex triplets (RMSD ranging from 0.634 Å to 0.881 Å for 124 atoms fit). This position in duplexes 1-3 is likely associated with greater deviations due to duplex end flexibility or crystal contact interactions, while deviations in duplex 4 are likely influenced by the structural changes induced by the adjacent d(TGA) triplet. Similarly low RMSD values are observed when comparing individual d(CGA) triplets from different duplexes (inter-duplex) (Figure S4). In the inter-duplex comparison, d(CGA) triplets from within duplex 3 exhibited the largest range of RMSD values when compared to triplets from other duplexes (0.459 Å to 1.043 Å for 124 atoms fit). Overall, the low RMSD values in the comparison of tandem and individual d(CGA) triplets illustrates that the ps-duplex form of d(CGA) repeat containing sequences are structurally isomorphous, even in different environmental contexts.

### d(CGA) helical and base-pair parameters

The high degree of similarity of the four unique ps-duplexes obtained from our crystal structures has allowed us to establish helical and base-pair parameters for this motif (Figure S5-S6). Like other d(CGA) and d(TGA) homo-duplex structures (36,37), the ps-duplexes form right-handed helices that lack distinct major and minor grooves. The d(CGA) ps-duplex form requires 9.0 ± 0.1 base pairs to complete one helical turn resulting in an average helical pitch of 32.2 Å ± 0.5 Å. The decreased helical pitch of the ps-duplex form, compared to B-DNA, is primarily a result of the large helical rise (5.0 Å ± 0.1 Å) and twist (84° ± 1°) associated with the inter-strand G/A base step. As previously observed (37), there is a notable difference in base pair parameters between purine and pyrimidines in the ps-duplex. Purine nucleotides adopt larger shear, propeller, stretch, and buckle angles to accommodate the hydrogen bonds while maintaining the duplex-stabilizing interstrand G/A base stacking interactions. We quantified the range of helical and base-pair parameters using 3DNA v2.4 (60) along each nucleotide position to highlight the periodic fluctuation of each parameter along individual d(CGA) triplets throughout the entire duplex. (Figure S5-S6).

We observed distinct ranges of phosphate backbone torsion angles (*α*, *β*, *γ*) among individual strands within each duplex (Figure S7). Notably, one strand in each duplex adopts a wide range of *α*, *β*, and *γ* torsion angles (246° ± 66°, 180° ± 36°, and 91° ± 66°, respectively), while the opposing strand of the duplex has a much narrower range (292° ± 5°, 164° ± 14°, and 59° ± 32°, respectively). The ability to adopt a wide range of torsion angles suggests that one strand is generally more flexible than its partner strand. This indicates that each strand within the ps-duplex is conformationally unique.

### Structural asymmetry is induced by the C-CH+ base pair

We assessed the overall symmetry of the ps-duplex form to determine how structural asymmetries are propagated through the ps-duplex. The linearity of each duplex along base pair units was measured by connecting the mid-point of the hydrogen bonding partners of each homo-base pair (Figure 4A). Each d(CGA) triplet exhibits a similar bending pattern centered around the largest deviation from linearity (25.0° ± 3.9°) at the C-CH+ base pair (Figure 4B). The G-G centric angle does not propagate significant deviations from linearity, while the magnitude of the A-A centric deviation is highly dependent upon the identity of the following nucleotide (C or T). When a C-CH+ base pair is present, the adjacent 5ʹ-A-A centric angle adopts a deviation (20.8° ± 4.3°) similar to the C-CH+ centric angle. Alternatively, the A-A centric deviation is smaller (9.0°) when followed by a T-T base pair (Figure 4B). Structural overlays of the A-(C/T) step indicate that this deviation coincides with the extension of one cytosine from the helical axis to align the Watson-Crick faces for the formation of the hemi-protonated C-CH+ base pair (Figure 4C), as previously noted (36). There is also a slight displacement of the adjacent adenosine on the same strand which could be required to accommodate the cytosine deviation. This contrasts with the T-T base pair which makes interactions in a perfectly symmetrical manner; therefore, the adjacent adenosine also remains unbent. We conclude that the A-A base pair provides structural flexibility to accommodate deviations from linearity induced by the C-CH+ base pair.

**Figure 4.**
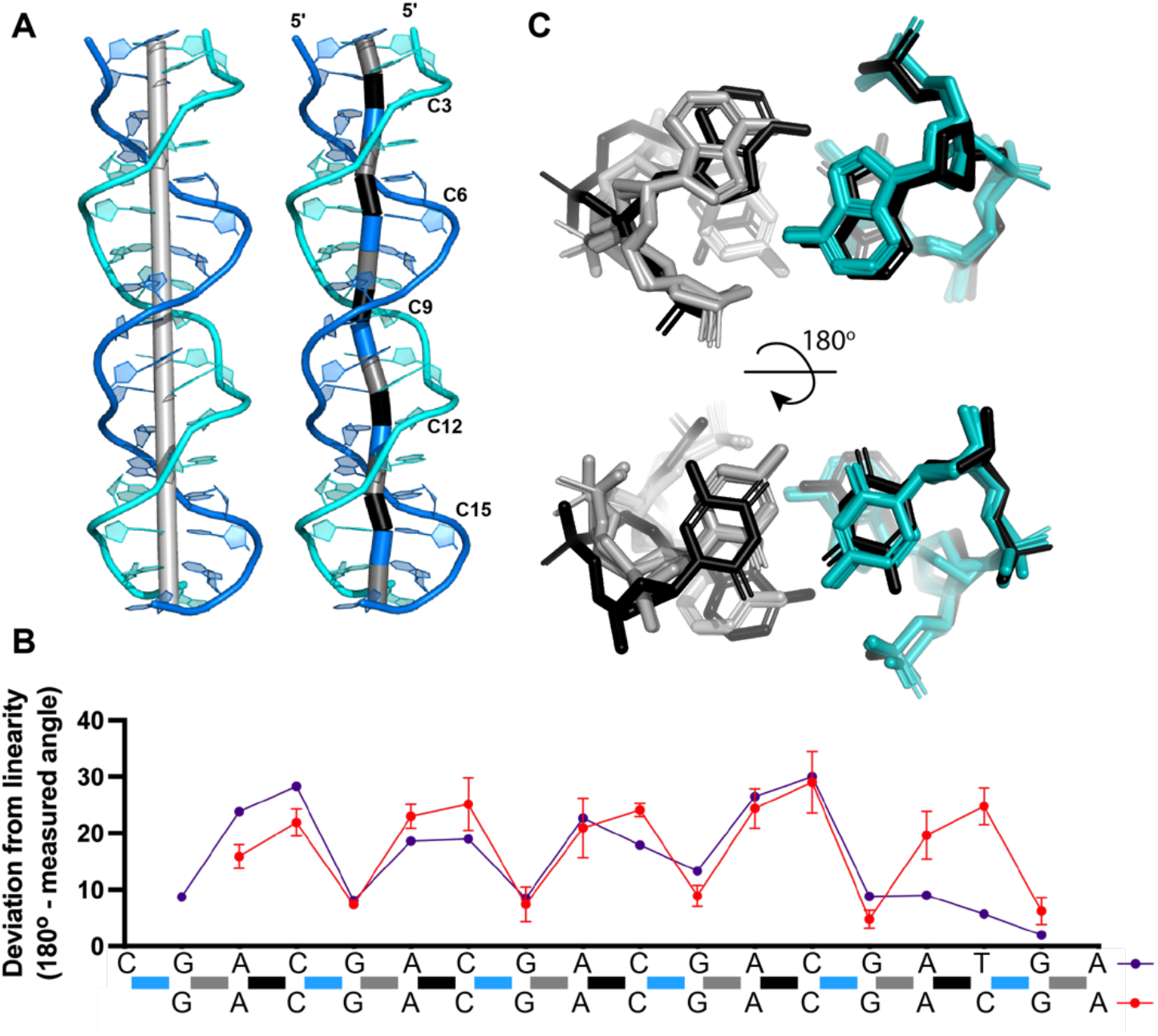
Parallel-stranded homo-duplex asymmetry. **A**. Deviations from linearity along base pairs of ps-duplex 1. (Left) Light gray cylinder represents the helical axis. (Right) The deviation from linearity of a base pair is measured as the angle between the two cylinders adjacent to the base pair of interest. Individual cylinders were created by connecting points placed at the midpoint of the hydrogen bonding partners of each base pair. The resulting cylinders are colored based on the identity of the base pairs they connect; G-A (gray), A-C (black), C-G (blue). **B**. Deviation from linearity of each base pair along the (CGA)_5_TGA (purple) or GA(CGA)_5_ (red) sequence. Colored bars along the sequence correspond to the same cylinders connecting base pairs from A. The angles measured for GA(CGA)_5_ are represented as the average of duplexes 1-3 and (CGA)_5_TGA is from duplex 4. **C**. Overlay of d(YGA) triplets within (CGA)_5_TGA, rotated 180° to highlight A-A and C-C base pairs. Five A/C steps (gray and teal) overlaid with one A/T step (black). Compared to the black strand, the teal strand does not show significant structural deviation, while the nucleotides within the gray strand are extended out of the helical axis.

### Each strand within the ps-duplex has unique structural character

Backbone torsion angle analysis and the duplex asymmetry suggested that the two strands of the ps-duplex have unique structural characteristics. These differences are correlated with two distinct hydrogen bond interactions that form within the A/C step between d(CGA) triplets (Figure 5A). The first hydrogen bond is between cytosine N4 (C-N4) and a non-bridging phosphate oxygen (O2P) of the previous adenosine within the same strand. There is no bond equivalent to the C-N4 to O2P bond in the T-T base pair, further suggesting that this bond could be influential in controlling the relative position of the C-CH+ and A-A base pairs. The second hydrogen bond is between the same non-bridging phosphate oxygen and the adenosine N6 (A-N6) of the opposing strand. Interestingly, depending on the strand within each duplex, there are unique differences in A-N6 to O2P and C-N4 to O2P bond lengths (Figure 5B). In one strand, herein referred to as “rigid,” all A-N6 to O2P and C-N4 to O2P bonds distances remain within 2.8 to 3.1 Å. However, within the opposite strand (referred to as “loose”) the same bond lengths increase beyond hydrogen bond distance. The average A-N6 to O2P and C-N4 to O2P distances within the loose strand are 4.1 ± 0.4 Å and 3.5 ± 0.1 Å, respectively. The cytosine that is displaced from the helical axis is always on the loose strand, where the increased bond lengths and wider range of torsion angles within the loose strand coincide with this displacement.

**Figure 5.**
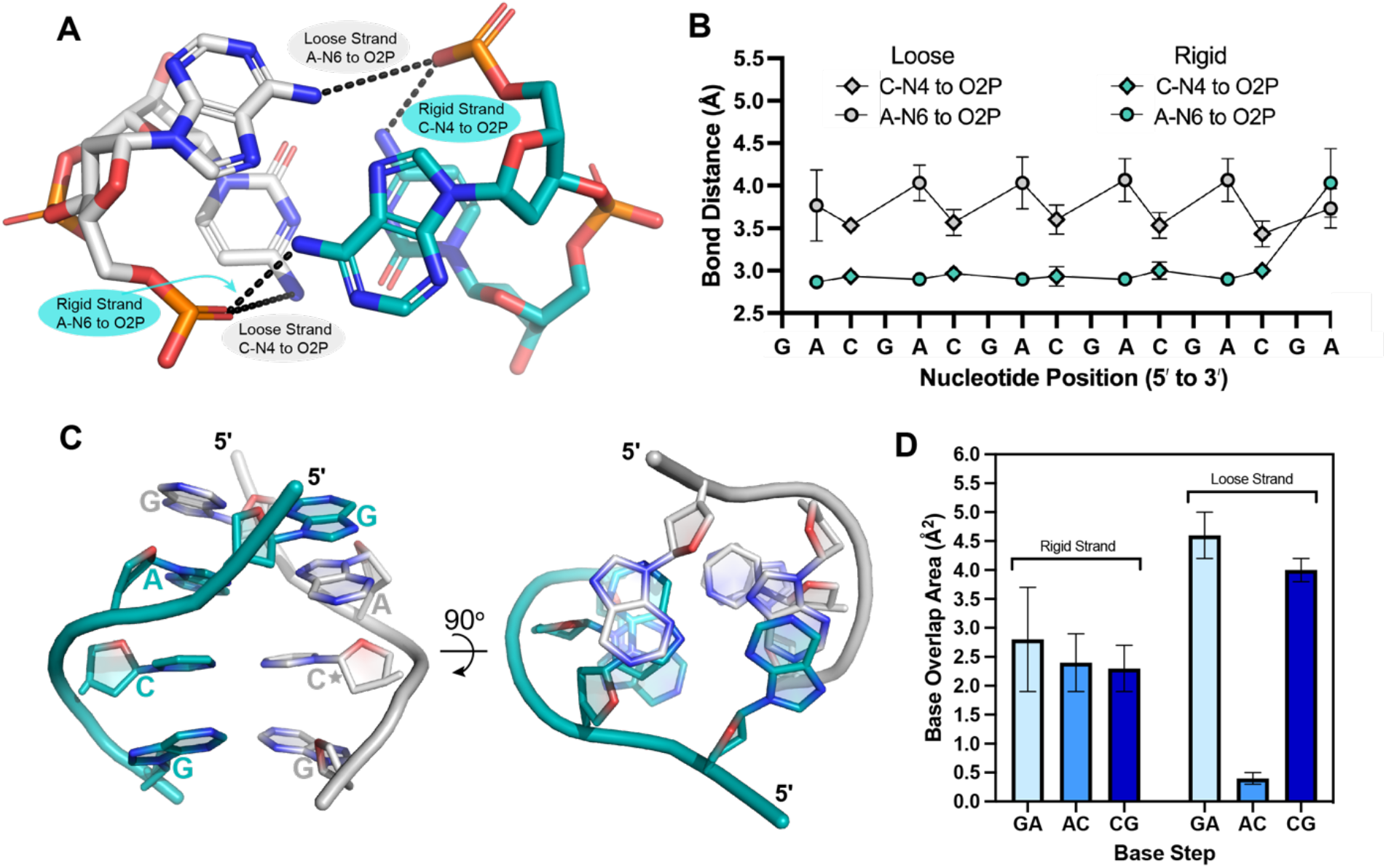
Hydrogen bond distances and base overlap areas are used to distinguish loose and rigid strands within the d(CGA) ps-duplex **A**. The A/C step highlighting the A-N6 to O2P and C-N4 to O2P interactions within loose (gray) and rigid (teal) strands. Chain A (duplexes 1 and 4), chain C (duplex 2), and chain E (duplex 3) have been characterized as loose strands. Chain B (duplexes 1 and 4), chain D (duplex 2), and chain F (duplex 3) have been characterized as rigid strands. **B**. Loose (gray) and rigid (teal) strand bond distances represented along the GA(CGA)_5_ sequence. A-N6 to O2P distances are plotted as circles and C-N4 to O2P distances are plotted as diamonds. Each data point represents the average distance measured from duplexes 1-3. Loose strand bond distances cycle between 3.5 ± 0.1 Å and 4.1 ± 0.1 Å depending on the identity of the nucleotide involved in the interaction while rigid strand bond distances remain between 2.8 to 3.1 Å, regardless of the interaction. **C**. Base overlap areas are different for loose and rigid strands. View of all unique base stacking interactions (inter-strand G/A, intra-strand A/C, and intra-strand C/G) that contribute to d(CGA) triplet stabilization. 90° rotation illustrates difference in stacking overlap area between strands. The rigid strand (teal) maintains consistent stacking overlap areas, while the loose strand (gray) is highly variable. The star denotes the cytosine that is extended from the helical axis. **D**. Base stack overlap areas are represented as averages of overlap areas from d(CGA) triplets from duplexes 1-4 and are shown for the respective base steps.

Accompanying the differences in hydrogen bonding are distinct differences in base stacking interactions between loose and rigid strands (Figure 5C). Base pair overlap areas (excluding exo-cyclic groups) calculated for each duplex using 3DNA v2.4 (60) indicate that intra-rigid strand A/C and C/G steps maintain similar overlap areas of 2.4 ± 0.5 Å^2^ and 2.3 ± 0.4 Å^2^, respectively (Figure 5D). The inter-strand G/A stacking interaction adjacent to the A/C step on the rigid strand also has a similar overlap area of 2.8 ± 0.9 Å^2^ (Figure 5D). However, the stacking areas of the loose strand are more variable. The A/C step on the loose strand has the lowest base overlap area (0.4 ± 0.1 Å^2^), while the G/A (inter-strand) and C/G (intra-strand) stacking interactions surrounding the A/C step have the highest stacking overlap area (4.6 ± 0.4 Å^2^ and 4.0 ± 0.2 Å^2^, respectively; Figure 5D). The large stacking interactions surrounding the bent A/C step within the loose strand contributes additional stabilization that may compensate for the increased base-to-phosphate hydrogen bond distances.

The overall structural asymmetry and accompanying differences in hydrogen bonding and base stacking interactions among strands are observed throughout each ps-duplex studied. Though it would be conceivable to expect the structural asymmetry to be propagated on a per-triplet basis, we observed the propagation on a per-strand basis over the entire length of the d(CGA) repeats. Thus, each ps-duplex is composed of two structurally unique strands where all triplets within a strand adopt either the loose or rigid character. The structural homogeneity of triplets within strands implies that duplexation of tandem d(CGA) triplets could occur in a cooperative manner. Further, the distinct conformations of each strand could play separate roles in accommodating the structural asymmetry. The rigid strand is the structural scaffold strand that maintains consistent hydrogen bonding and stacking interactions, while the loose strand provides structural flexibility to stabilize and accommodate deviations from linearity induced by the C-CH+ base pair.

### d(YGA) triplets are structurally compatible, but not identical

It has been hypothesized that d(TGA) triplets could be useful discriminators in the programmable pairing of long stretches of d(CGA) triplets based on slight structural and thermodynamic deviations incurred by the 5ʹ-nucleotide (31). Previous crystal structures have reported differences in d(CGA) (36) and d(TGA) (37) triplets from separate sequence contexts but have not yet examined the structural compatibility when d(YGA) triplets are present within the same sequence. The crystal structure of (CGA)_5_TGA has allowed us to evaluate the structural compatibility of d(CGA) and d(TGA) triplets within the same DNA sequence. We observed that the incorporation of a 3ʹ-d(TGA) triplet significantly alters the bond distances of loose and rigid strands in upstream d(CGA) triplets. Within the d(CGA) triplet directly adjacent to the d(TGA) triplet, the rigid strand C-N4 to OP2 and A-N6 to OP2 bond distances increase from an average of 3.0 Å to 4.4 Å and 4.9 Å, respectively, while the loose strand A-N6 to OP2 distance decreases from 4.0 Å to 3.5 Å (Figure S8A). The C1ʹ-C1ʹ distance for the T-T homo base pair is 1.4 Å wider than the C-CH+ homo-base pair, therefore, upstream swelling of the rigid strand could be required to accommodate the wider T-T homo-base pair (Figure S8B). Increased base overlap areas of the G/A steps adjacent to the d(TGA) triplet could also contribute additional stabilization to compensate for the extended rigid strand bond distances (Figure S8C,D). Enthalpic destabilization observed in sequences containing d(TGA) triplets was previously attributed to the loss of one hydrogen bond from replacing the hemi-protonated C-CH+ with a T-T base pair (31). The structure described here further suggests this destabilization could also be due to the loss of the C-N4 to O2P hydrogen bond and swelling of adjacent d(CGA) triplets that coincide with the addition of a T-T base pair. The incorporation of a d(TGA) triplet at the 3ʹ-end of a long stretch of d(CGA) triplets does not disrupt the overall ps-duplex structure but induces slight structural changes in the adjacent d(CGA) triplet.

Though d(CGA) and d(TGA) triplets are not structurally identical within the ps-duplex, they could be used to control rigid and loose strands. Interestingly, U^Br^GA triplets have been shown to offer increased stability to the ps-duplex via the formation of a halogen bond with the phosphate oxygen of an adjacent adenosine (37). This indicates the valuable prospective of the U^Br^GA triplet in the rational design of d(CGA) containing ps-duplexes. To fully evaluate the potential use as discriminator triplets, structural analysis of d(CGA) repeat sequences containing internal d(TGA) triplets (and U^Br^GA triplets) are needed to understand the effect of d(TGA) incorporation on downstream d(CGA) triplets.

### Prospects for d(CGA)-based ps-duplexes in technology development and biology

The crystal structures described here have allowed us to characterize the d(CGA) triplet repeat motif in the ps-duplex form and establish structural features for its use as a building block in DNA nanotechnology applications. The generalized helical and base parameters established by these structures will serve as constraints for the incorporation of d(CGA)-based triplets into rational structure design. Particularly, requiring an integer number of base pairs per turn (9.0 ± 0.1 base pair) simplifies its use from a design perspective, as incorporation of three d(CGA) repeats completes exactly one helical turn. Consistent with previous d(CGA) base-paired triplets, we observed a structural asymmetry that is propagated throughout each duplex. Our crystal structures containing multiple tandem d(CGA) triplets have demonstrated that this asymmetry is propagated on a per-strand basis, where all triplets within each strand adopt a specific conformation and play a unique role in accommodating the asymmetry. Specifically, the rigid strand serves as the structural scaffold that maintains hydrogen bonding and stacking interactions, while the loose strand provides structural flexibility to stabilize and accommodate deviations from linearity induced by the C-CH+ base pair. The distinct structural character between triplets within rigid and loose strands could be a useful tool in the structural programming of DNA based architectures where specific control of each strand within the duplex may be desirable. The structural similarity of tandem and individual d(CGA) base-paired triplets obtained from different solution environments demonstrates that the ps-duplex form is a robust and highly predictable structure. This strongly suggests that the d(CGA) motif can be used to reliably integrate the ps-duplex form into nanostructures. This motif has the added benefit of allowing conditional control of the ps-duplex form in solution through mild fluctuations in pH or the addition of crowding agents.

Though the formation of stable alternative DNA structures may be desirable for the rational design of DNA-based architectures, they may be unfavorable or selected against in biological systems. Specifically, repeat sequences that form alternative structures could undergo greater instability due to the challenges they present to replication machinery (46). However, there is also the possibility that readily formed, thermodynamically stable structures would be selected against evolutionarily, as they may have an even greater impact on endogenous replication systems. Interestingly, d(CGA) triplet repeat expansions have not been implicated in human disease and are found least frequently in eukaryotic genomes (42,43). This raises the interesting possibility that their propensity to form highly stable ps-duplex structures could be a significant factor contributing to the under-representation of the d(CGA) triplet repeat in eukaryotic genomes. Though questions have arisen about the likelihood of C-CH+-dependent structures forming in vivo, mounting evidence – including data presented here – suggest that such structures can form at near-neutral pH under crowding conditions.

## DATA AVAILABILITY

Atomic coordinates and structure factors for the reported crystal structures have been deposited in the Protein Data Bank under the accession codes 7SB8 and 7SB9.

## SUPPLEMENTARY DATA

Supplementary Data are available at NAR online.

## ACKNOWLEDGEMENT

This work is based upon research conducted at the Northeastern Collaborative Access Team beamlines, which are funded by the National Institute of General Medical Sciences from the National Institutes of Health (P30 GM124165). The Eiger 16M detector on 24-ID-E is funded by a NIH-ORIP HEI grant (S10OD021527). This research used resources of the Advanced Photon Source, a U.S. Department of Energy (DOE) Office of Science User Facility operated for the DOE Office of Science by Argonne National Laboratory under Contract No. DE-AC02-06CH11357.

