## Supplemental Information for "The parallel-stranded d(CGA) duplex is a highly predictable structural motif with two conformationally distinct strands"

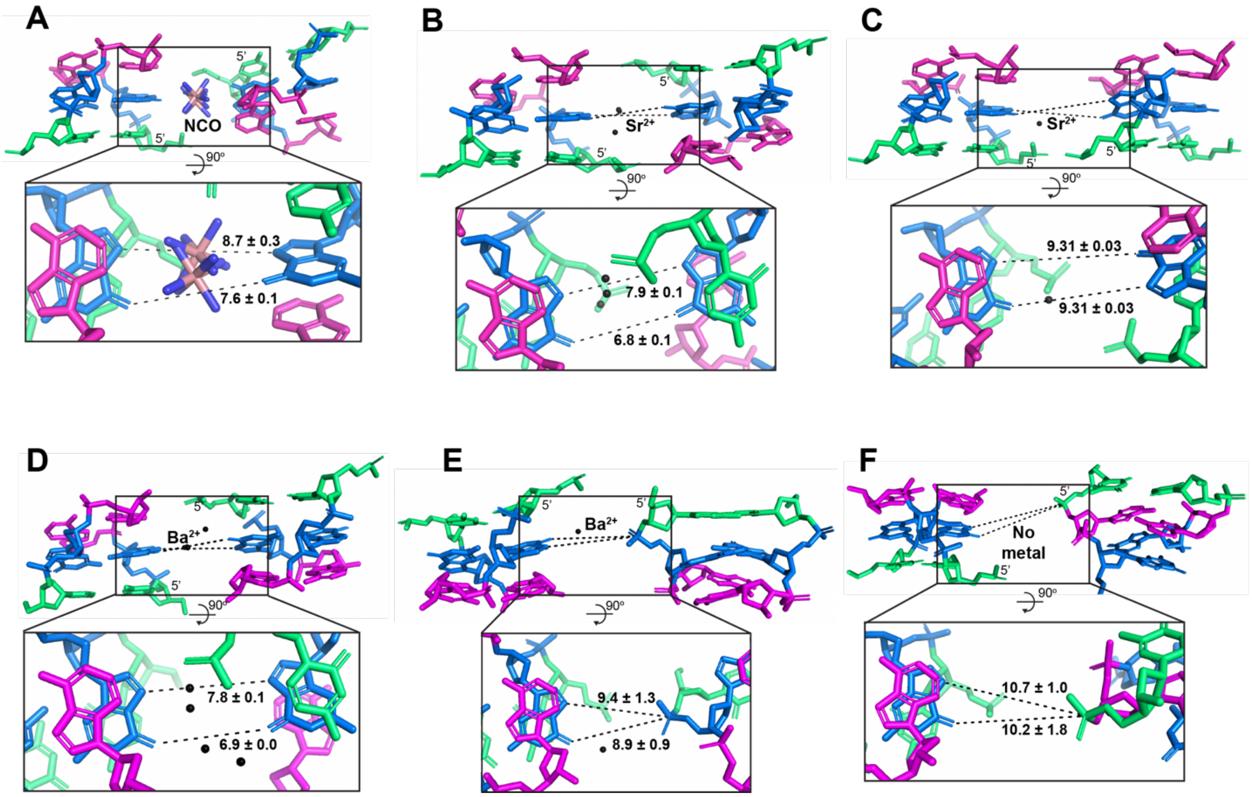


**Figure S1.** Unique cation mediated crystal packing interactions were observed within GA(CGA)$_5$ (**A-C**) and (CGA)$_5$TGA (**D-F**) structures. Cations (NCO, Sr$^{2+}$, and Ba$^{2+}$) are always found positioned between two guanosines from separate duplexes. d(CGA) triplets surrounding each cation are shown and colored as follows: cytosine (green), guanosine (blue), adenosine (magenta). The averaged bond distances between cation mediated guanosines are shown. Although there are unique packing arrangements found in each dataset, the resulting ps-duplex structure is unaltered.

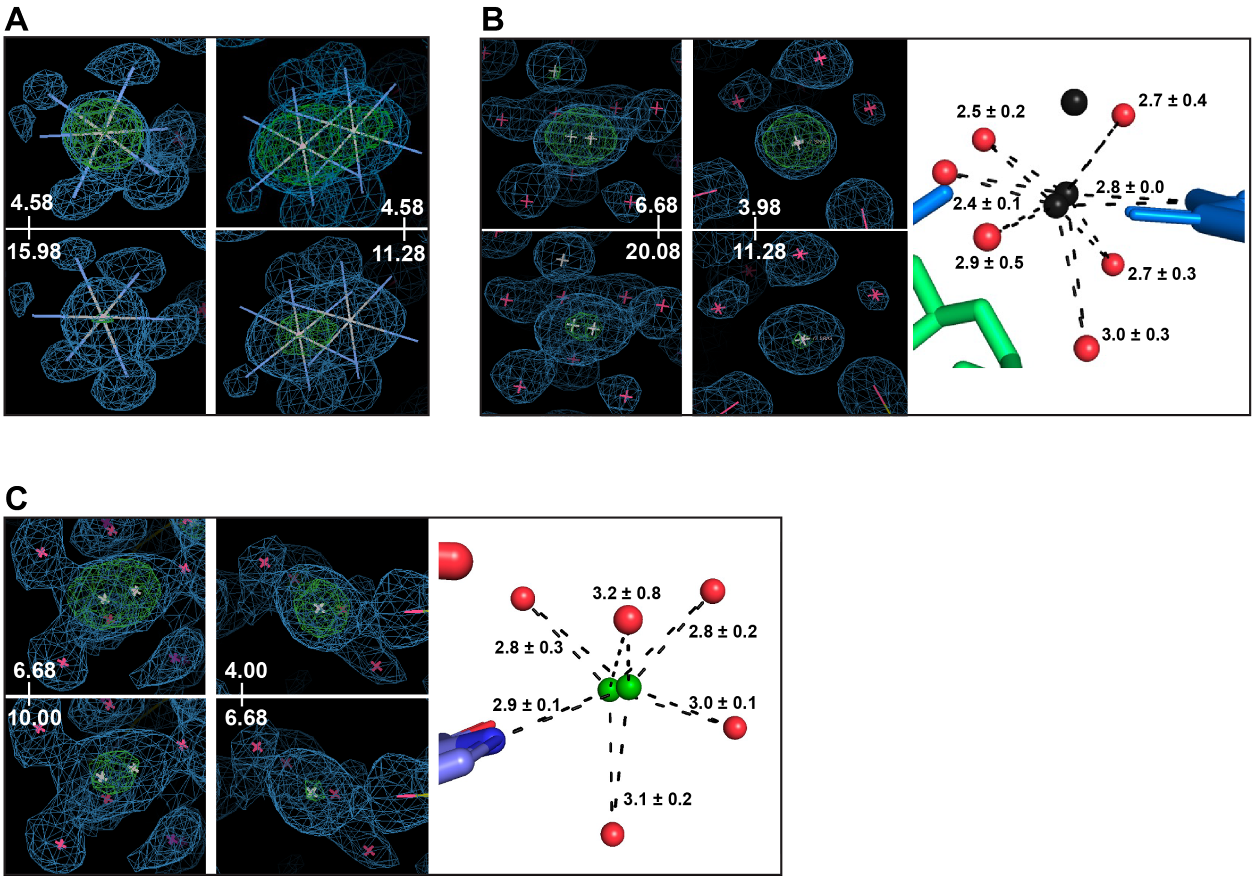


**Figure S2.** Anomalous difference maps and coordination distances used to verify the placement of cations **A.** NCO **B.** $Sr^{2+}$ **C.** $Ba^{2+}$. Anomalous difference maps are shown at representative high and low $\sigma$ values for each cation. The $2mF_o$-$DF_c$ electron density map is contoured to 1 $\sigma$. Average coordination distances are shown for $Sr^{2+}$ and $Ba^{2+}$.

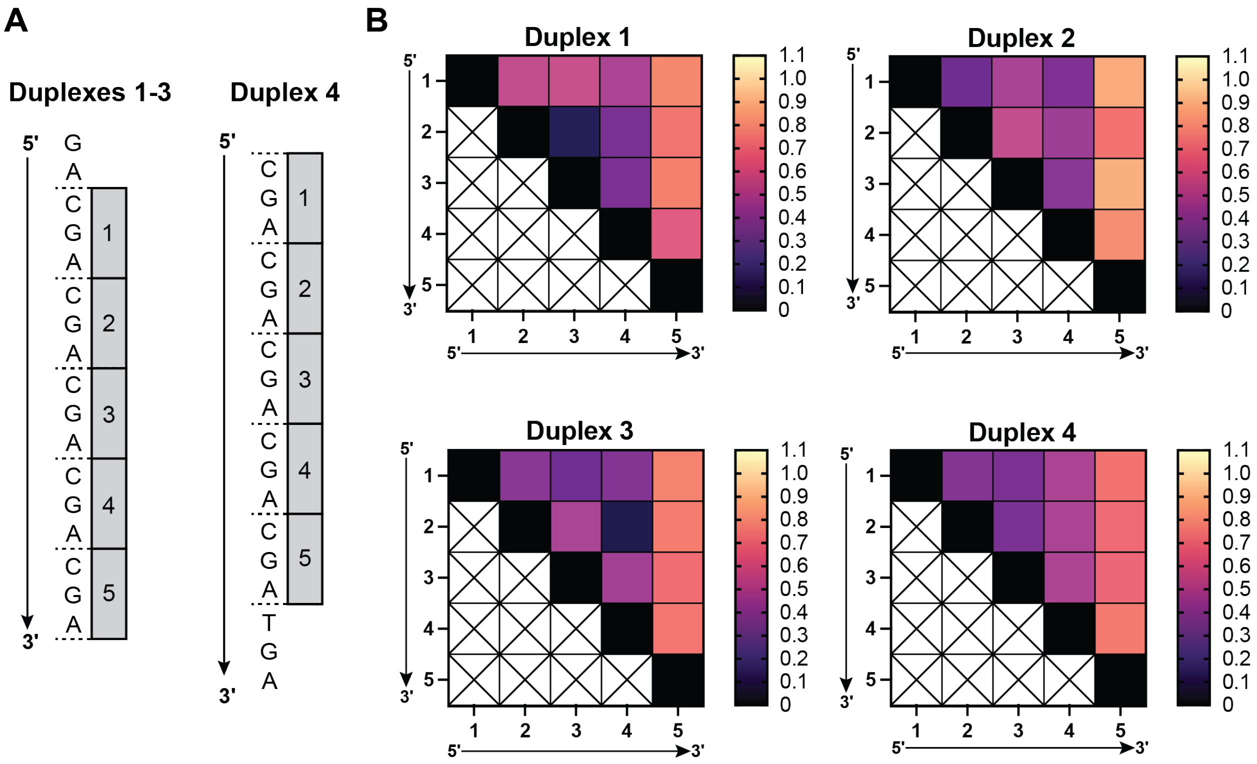


**Figure S3.** Individual intra-duplex d(CGA) triplets are highly uniform. **A.** Individual d(CGA) triplets are numbered 1-5 starting at the first complete d(CGA) triplet within each duplex. Specific duplex numbering is shown for each duplex. **B.** RMSD values obtained from the alignment of d(CGA) triplets within the same duplex (intra-duplex). Alignments containing the 3' d(CGA) triplet (position 5) from each duplex always result in the highest RMSD value, indicating that this position is associated with structural flexibility.

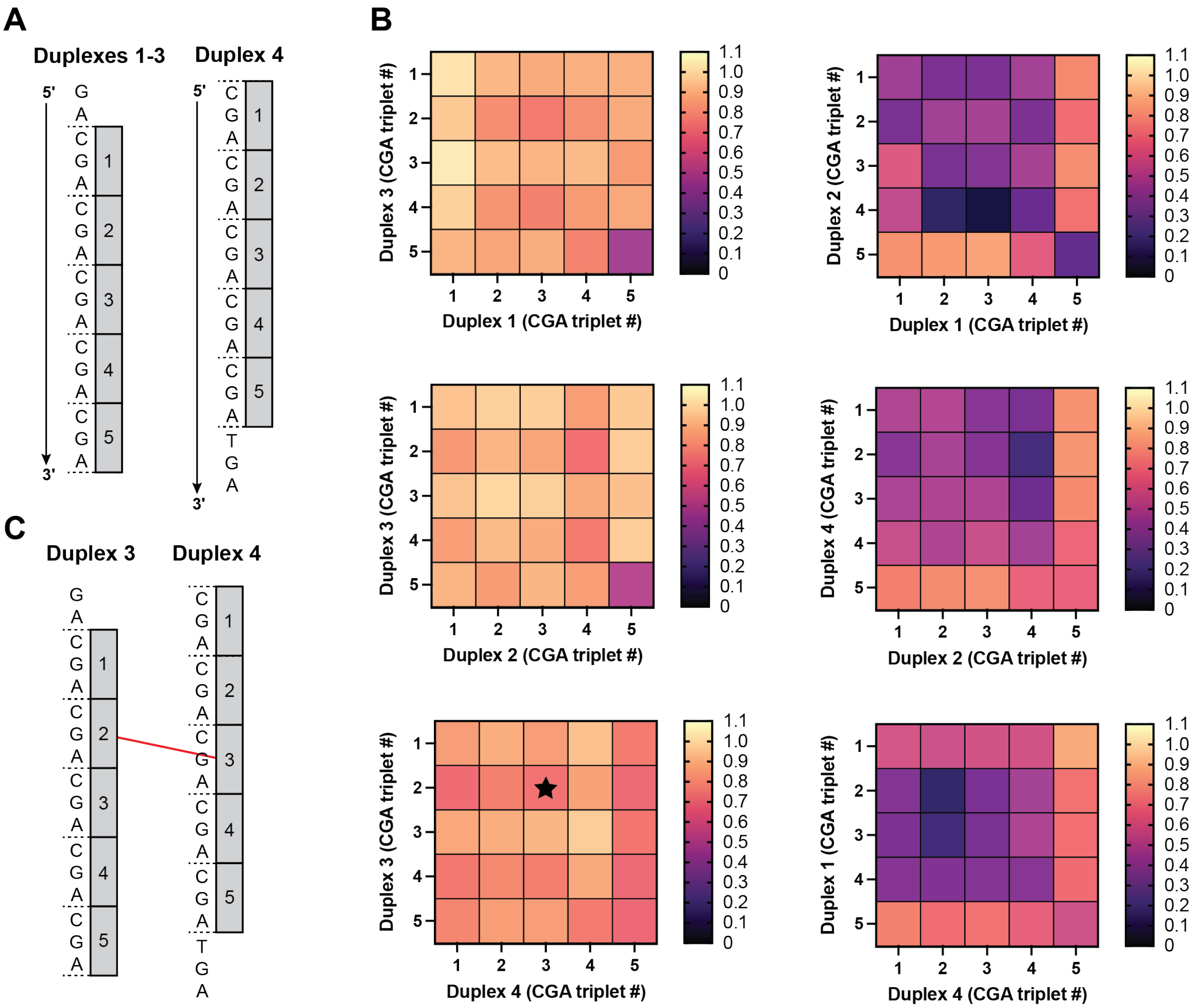


**Figure S4.** Individual inter-duplex d(CGA) triplets are highly uniform. **A.** Individual d(CGA) triplets are numbered 1-5 starting at the first complete d(CGA) triplet within each duplex. Specific duplex numbering is shown for each duplex. **B.** RMSD values obtained from the alignment of d(CGA) triplets from different duplexes (inter-duplex). d(CGA) triplets from duplex 3 have the highest RMSD values. **C.** Schematic representation of one set of d(CGA) triplets compared within duplexes 3 and 4 indicated by the star in B.

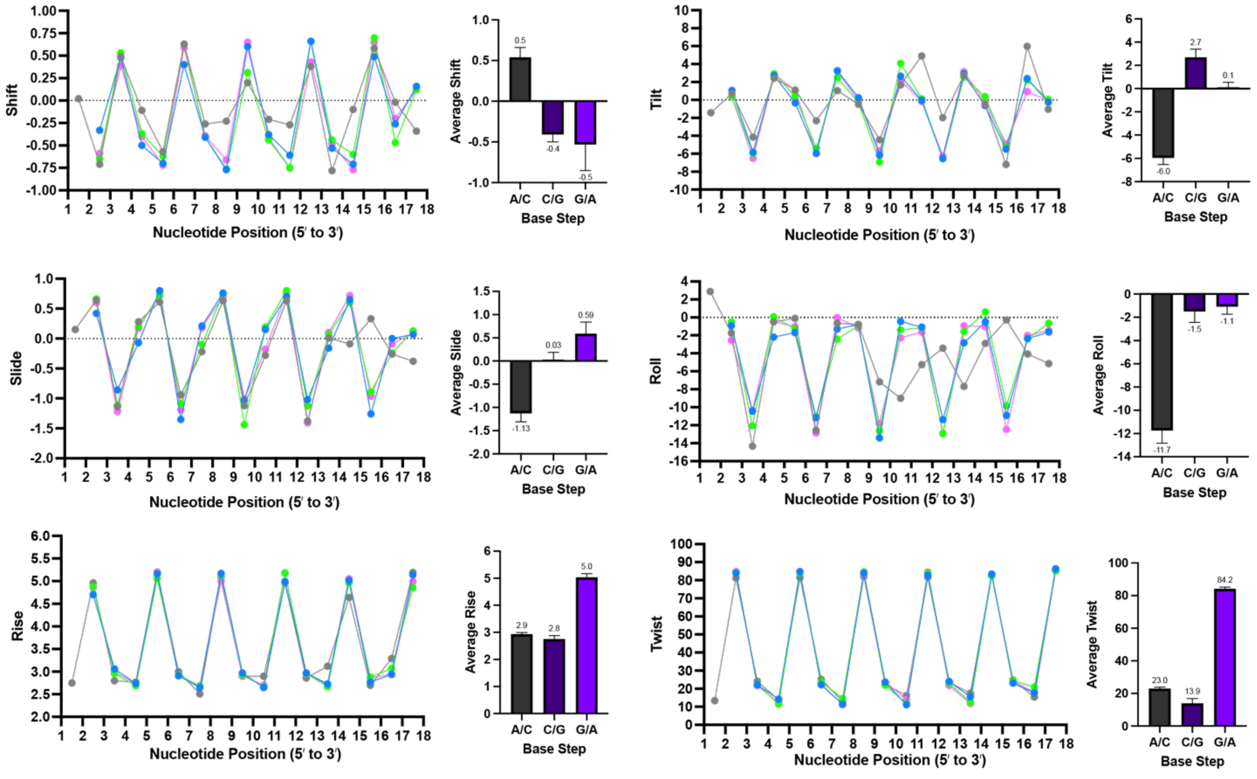


| | |
|---|---|
| **Duplex 4** | **C G A C G A C G A C G A C G A T G A** |
| Nucleotide position | 1 2 3 4 5 6 7 8 9 10 11 12 13 14 15 16 17 18 |
| **Duplexes 1-3** | **G A C G A C G A C G A C G A C G A** |

**Figure S5.** Helical parameters were obtained from 3DNA (60) and are shown for each base step along duplexes 1 (blue), 2 (green), 3 (pink), and 4 (gray). Bar graphs represent the average parameter for each respective step in duplexes 1-4.

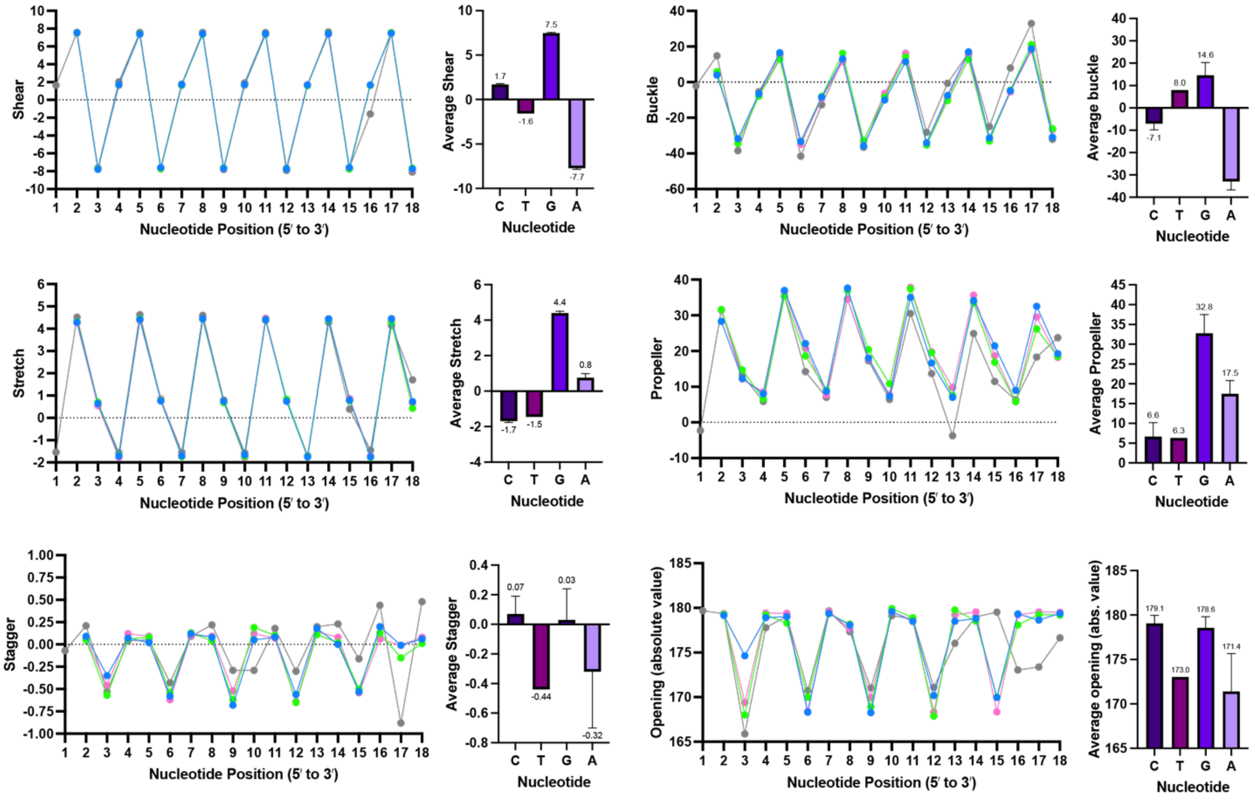


**Duplex 4** C G A C G A C G A C G A C G A T G A

Nucleotide position 1 2 3 4 5 6 7 8 9 10 11 12 13 14 15 16 17 18

**Duplexes 1-3** G A C G A C G A C G A C G A C G A

**Figure S6.** Simple base-pair parameters were obtained from 3DNA (60) and are shown for each base pair along duplexes 1 (blue), 2 (green), 3 (pink), and 4 (gray). Bar graphs represent the average parameter for each respective base pair in duplexes 1-4.

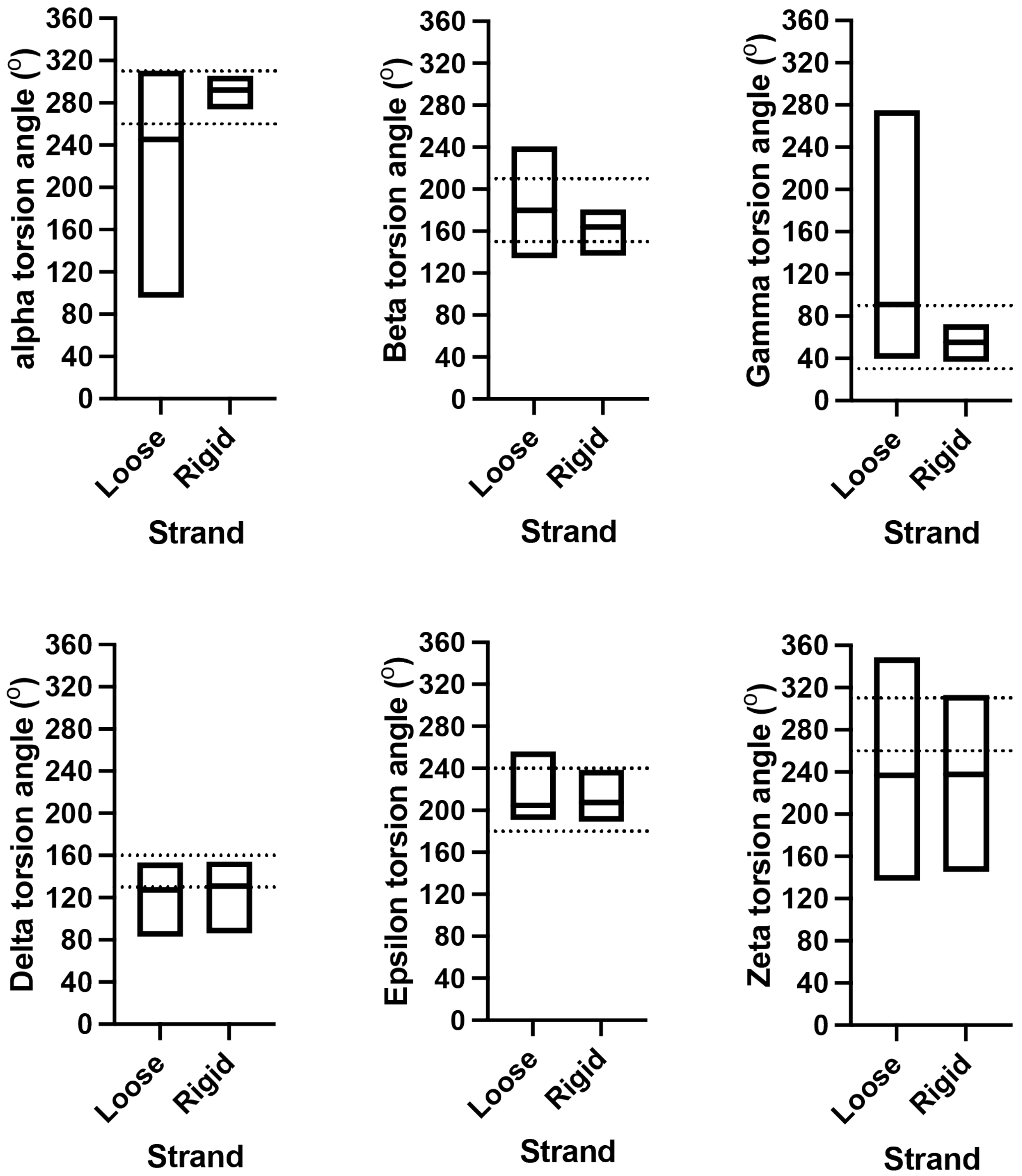


**Figure S7.** Torsion angles (α, $\beta$, $\gamma$, $\delta$, $\varepsilon$, and $\zeta$) obtained from 3DNA (60). Torsion angles were calculated for each strand from all nucleotides in duplexes 1-4, excluding the TGA triplet. There is a difference in the range of accepted α, $\beta$, and $\gamma$ angles depending on the strand, indicating that each strand has unique structural character. Plots illustrate the average and standard deviation of each angle. Dashed lines represent the torsion angles observed for ps-duplexes in the literature (61).

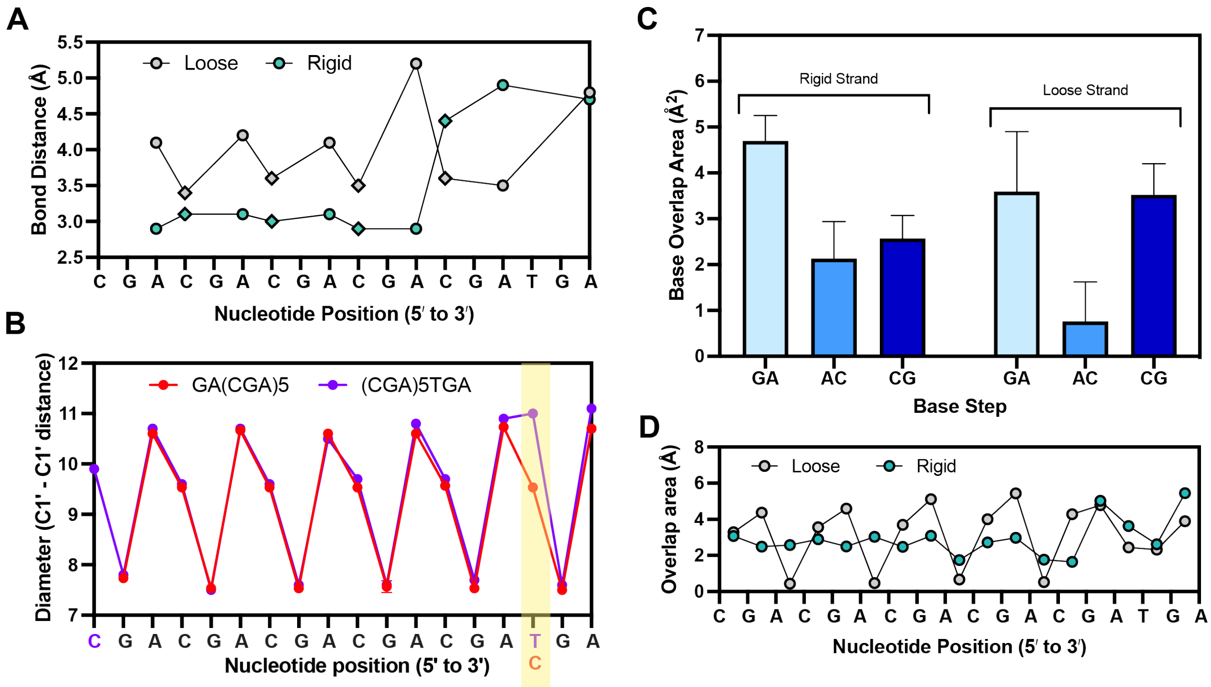


**Figure S8.** Loose and rigid strand bond distances and base overlap areas for $(CGA)_5$TGA. The 3'-d(TGA) triplet disrupts adjacent d(CGA) nucleotide-backbone hydrogen bonds and base stacking interactions. **A.** Loose (gray) and rigid (teal) strand nucleotide to backbone bond distances plotted along the $(CGA)_5$TGA sequence. A-N6 to O2P distances are plotted as circles and C-N4 to O2P distances are plotted as diamonds. The rigid strand bond distances significantly increase in the d(CGA) triplet directly upstream of the d(TGA) triplet. **B.** Duplex diameter as measured by the C1' to C1' distance of each base pair along sequences GA$(CGA)_5$ (red) and $(CGA)_5$TGA (purple). GA$(CGA)_5$ data points are represented as average measurements from duplexes 1-3. The difference in diameter between C-C and T-T base pairs is highlighted in yellow. **C.** Base stack overlap areas are represented as averages of overlap areas from d(CGA) triplets in duplex 4 and are shown for each respective base step. **D.** Intra-strand (A/C or C/G) and inter-strand base step overlap areas plotted for each step along the $(CGA)_5$TGA sequence. Teal points represent base overlaps between nucleotides in rigid strands and gray points represent base overlap between nucleotides in loose strands. In the (G/A) inter-strand base step, teal points represent the base overlap of a rigid strand guanosine on top of a loose strand adenosine in the 5' to 3' direction. Gray points represent the reverse; a loose strand guanosine stacking on top of a rigid strand adenosine in the 5' to 3' direction.
